# tRNA-derived fragments are elevated in the aging brain and may contribute to neurodegeneration

**DOI:** 10.64898/2026.05.15.725455

**Authors:** Lien D. Nguyen, Ami Kobayashi, Alain Ndayisaba, Vikram Khurana, Pavel Ivanov, Anna M. Krichevsky

## Abstract

tRNA-derived fragments (tRFs) are a class of small noncoding RNAs with emerging roles in stress responses and gene regulation, yet their dynamics across tissues and brain regions during aging remain poorly understood. Here, we systematically profiled age-associated changes in tRFs using three independent mouse small RNA-seq datasets spanning multiple organs and brain regions. Nuclear-encoded tRFs were the only small RNA class showing a strong, progressive accumulation with age, a pattern that was specific to the brain and broadly distributed across brain regions. Fragment length distributions, boundary profiles, and coverage maps were consistent with amplified cleavage at conserved sites, implicating angiogenin as a primary driver. Age-associated increases in specific tRF species, including 5’Cys^GCA^, 5’Glu^CTC^, and 5’Gly^GCC^, were validated by northern blotting and RT-qPCR. Analogous upregulation was observed in human frontal lobe tissue from frontotemporal dementia patients and in cerebrospinal fluid from traumatic brain injury patients, suggesting that tRF accumulation is further amplified under neurological stress. Together, these findings establish nuclear-encoded tRFs as a small RNA class that accumulates selectively in the aging brain, with potential roles in neurodegeneration, and as biomarkers and targets for therapeutic intervention.

## INTRODUCTION

Aging is a progressive biological process characterized by the gradual decline of cellular and tissue function and an increased risk of death over time. Although aging itself is not classified as a disease, it is a major contributor to many disorders and is the strongest risk factor for most neurodegenerative diseases, including Alzheimer’s disease (AD) and Parkinson’s disease (PD)^1^. The brain is particularly vulnerable to age-related molecular changes, especially due to the post-mitotic nature of neurons, their high metabolic demands, and their limited capacity for self-renewal^1^. The molecular mechanisms underlying brain aging and neurodegeneration have been extensively investigated, with particular focus on protein aggregates such as β-amyloid and Tau in AD, and αSynuclein in PD^2^. Recently, small non-coding RNAs have emerged as important modulators of gene expression, cellular stress responses, and neuronal homeostasis^3^. However, most studies investigating small RNAs in aging and neurodegeneration have focused primarily on microRNAs (miRNAs), leaving other classes of small RNAs relatively understudied.

Transfer RNAs (tRNAs), the most abundant class of RNA in the cell by molar count^4^, are traditionally known for their canonical role in translation, where they deliver amino acids to the ribosome for protein synthesis. Under stress conditions, tRNAs can be cleaved by nucleases, including angiogenin^5,6^, to generate tRNA-derived fragments (tRFs) with diverse regulatory functions^4^. This cleavage is an evolutionarily conserved stress response observed across species, and in humans, approximately 500 nuclear-encoded tRNA genes and 22 mitochondrial tRNA genes can be cleaved to produce hundreds to thousands of unique small RNA fragments^7,8^. tRFs have been shown to inhibit translation^5,9,10^, compete with messenger RNA (mRNA) for ribosome binding^11^, participate in miRNA-like RNA silencing pathways^12^, and modulate various cellular processes such as autophagy^13^ and proliferation^14^. Recent studies indicate that specific tRFs are elevated in the aging mouse brain^15^, as well as in brains and biofluids from AD^16,17^ and PD^18,19^ patients. Consistent with these observations, we recently identified stress-responsive tRFs that directly bind to and promote tau phosphorylation and aggregation in cellular and animal models of AD^20^. These findings suggest that tRFs may contribute to pathological processes in brain aging and neurodegeneration. Furthermore a previous study observed a monotonous increase in 3’ tRFs and a less consistent increase in 5’ tRFs with age in the rat brain, but were limited to three brains each across three time points^21^. Despite these advances, the dynamics, tissue specificity, and molecular properties of tRFs across mammalian tissues and brain regions during aging remain poorly defined.

To address this gap, we systematically profiled age-related changes in tRFs using publicly available small RNA-seq datasets spanning multiple mouse organs^22^ and brain regions^23^ across aging. This analysis revealed that nuclear-encoded tRFs are the small RNA class most strongly and selectively elevated with age in the brain and broadly distributed across brain regions. Age-associated increases were driven by amplified cleavage at conserved, structurally defined sites rather than nonspecific RNA degradation, consistent with a role for the RNase angiogenin. Increases in specific tRF species, including 5’Cys^GCA^, 5’Glu^CTC^, and 5’Gly^GCC^, were validated by northern blot and RT-qPCR across multiple brain regions. Analogous tRF upregulation was observed in human frontotemporal dementia (FTD) brain tissue and in cerebrospinal fluid from TBI patients, suggesting that the age-associated tRF response is further amplified under neurological stress. Together, these findings indicate that specific tRFs accumulate selectively in the aging brain and may contribute to pathological processes linked to neurodegeneration, highlighting them as potential biomarkers and therapeutic targets.

## RESULTS

### Nuclear-encoded tRNA-derived fragments accumulate with age specifically in the brain

To determine the dynamics of small RNA changes with age, we first reanalyzed a publicly available small RNA-seq dataset (GSE111279) comprising bulk small RNA profiles from the right brain hemisphere of male C57BL/6 mice across five age groups (n = 5 per group). Sequencing reads were processed using our small RNA analysis pipeline involving sequential alignment to reference RNA classes (Sup. Fig. 1A), which achieved greater than 95 percent overall mapping efficiency across samples (Sup. Fig. 1B). tRNA-derived reads were further classified into 5′ tRFs, 3′ tRFs, internal tRFs, or full-length tRNAs based on their alignment positions relative to annotated tRNA genes. Given the maximum read length of 50 nt in this small RNA-seq library (Sup. Table 1), full-length tRNA reads were negligible. Because many tRNA isodecoders differ by only one or two nucleotides, tRNA-derived reads frequently exhibit high multimapping rates. Accordingly, isodecoders within each tRF family showed highly correlated expression patterns across samples (median Spearman ρ = 0.8 with family sum; ∼90% tRFs showed a concordant direction of association with age relative to their family sum, Sup. Fig. 1C-D) and were therefore collapsed into single family-level features for downstream analyses, reducing ∼224 isodecoders to 51 entries per tRF class.

Nuclear-encoded tRFs were the only RNA classes showing a strong age-associated increase in the brain (Fig. 1A). Among nuclear tRF species with significant age associations, positive correlations predominated across all three fragment classes: 49% of 3′ tRFs, 32.7% of internal tRFs, and 25% of 5′ tRFs were significantly positively correlated with age. By contrast, mitochondrial tRFs (Mt-tRFs) showed predominantly negative age associations. Other small RNA classes examined, including miRNAs, piRNAs, scaRNAs, snoRNAs, and snRNAs, had significant positive age correlations in fewer than 11% of species (Fig. 1A). miRNAs and scaRNA additionally showed strong negative age association, though this was not replicated in another dataset (Sup. Fig. 2A-B). The increase in nuclear tRFs was also evident at the level of median class abundance, with nuclear tRFs showing a progressive rise across age groups, particularly between 15 and 24 months, while other RNA classes remained relatively stable (Fig. 1B). Principal component analysis (PCA) of normalized nuclear tRF expression revealed a clear age-dependent trajectory along PC1 (27.1% variance), with younger (2, 9, and 15 mo) and older (24 and 30 mo) brains segregating toward opposite poles of the axis (Fig. 1C)

**Figure 1.**
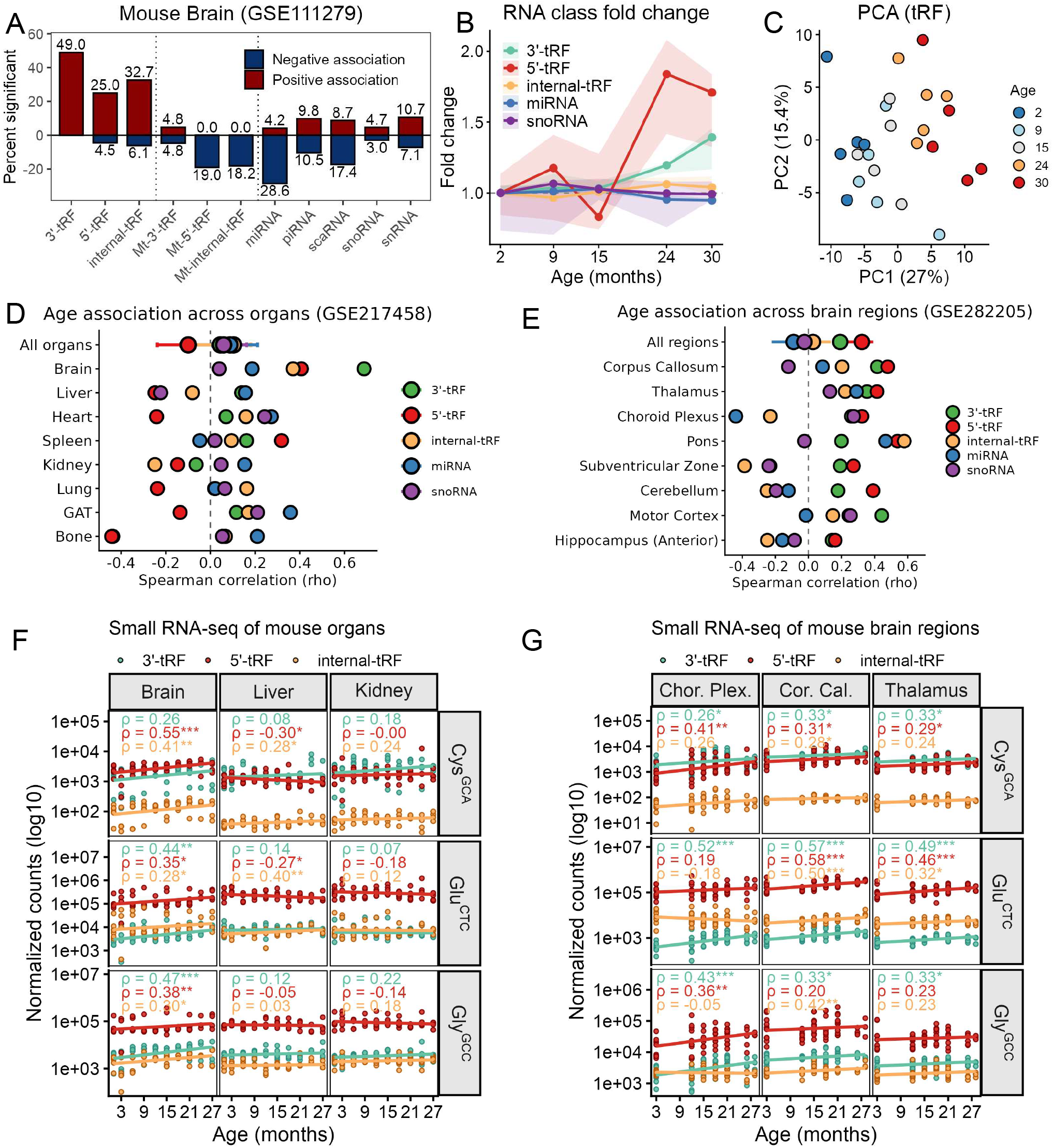
Nuclear-encoded tRNA-derived fragments (tRFs) show significant elevation with age specifically in the mouse brain. **(A)** Percentage of RNAs within each biotype with a significant positive (red) or negative (blue) Spearman correlation with age in mouse brain small RNA-seq data (GSE111279; 5 age groups, n = 5 male mice per group). Significance threshold: FDR-adjusted p < 0.05 (Benjamini-Hochberg). **(B)** Fold change in abundance of major RNA classes relative to the youngest age group, computed from DESeq2-normalized counts. Lines represent median fold change across all RNAs within each class per age group; shaded regions denote the interquartile range (IQR). **(C)** Principal component analysis (PCA) of DESeq2-normalized nuclear-encoded tRF expression across samples. **(D)** Spearman correlation (ρ) between RNA abundance and age across organs (GSE217458; 16 organs total, 8 shown; 10 age groups, n = 4-6 male and female mice per group). Each point represents the median ρ across all RNAs within a biotype per organ; error bars indicate IQR. **(E)** Spearman correlation (ρ) between RNA abundance and age across brain regions (GSE282205; 15 regions total, 8 shown; 8 age groups, n = 6-10 male and female mice per group). Each point represents the median ρ across all RNAs within a biotype per region; error bars indicate IQR. **(F-G)** DESeq2-normalized expression of Cys^GCA^, Glu^CTC^, and Gly^GCC^ tRF families across age, stratified by tissue:whole organs and (F) selected brain regions. Points represent individual samples; lines indicate linear regression fits, shown separately for each fragment class (3′, 5′, and internal tRFs). Spearman correlation coefficients (ρ) and unadjusted p-value significance levels are shown within each panel (*p < 0.05, **p < 0.01,***p < 0.001). y-axes show log10-scaled normalized counts.

To assess organ and regional specificity, we extended our analysis to two additional publicly available datasets: a multi-organ aging dataset (GSE217458)^22^ encompassing 16 mouse organs across ten age points (n = 4 to 6 male and female mice per group), and a brain region-resolved aging dataset (GSE282205)^23^ encompassing 15 regions across eight age points (n = 6 to 10 male and female mice per group). Interestingly, while these studies focused primarily on miRNAs, both noted an age-associated increase in tRNA-derived reads in the brain and brain regions. Consistent with findings in GSE111279, nuclear-encoded tRFs were, again, the only RNA class showing positive age correlations in the brain, with minimal changes in other RNA classes (Sup. Fig. 2A-B). Comparison of age-correlated tRF accumulation across 16 organs demonstrated that this positive correlation was largely restricted to the brain, with other tissues showing near-zero or negative median correlations (Fig. 1D, Sup. Figure 2C). Analysis across 15 brain regions revealed that positive age-associated tRF accumulation was observed in at least 8 of the 15 brain regions, and correlations were strongest in the corpus callosum, thalamus, and choroid plexus (Fig. 1E, Sup. Figure 2D-F).

At the level of individual tRNA families, representative nuclear-encoded tRFs including those derived from Cys^GCA^, Glu^CTC^, and Gly^GCC^ showed significant increases with age in the brain and various brain regions, while non-brain tissues showed no consistent directional trend across fragment types from the same family (Fig. 1F-G). Multiple fragment types derived from the same parental tRNA showed coordinated age-associated trajectories, suggesting increased cleavage of specific nuclear tRNAs with age in the brain. Together, findings from three independent datasets consistently support that nuclear-encoded tRFs are uniquely and selectively elevated during brain aging in mice.

### tRNA-derived fragment abundance increases with age at conserved cleavage sites

To determine whether the age-associated rise in tRF levels reflects increased cleavage at specific sites, or a global, non-specific accumulation of fragments, or technical artefacts arising from age-associated alterations in tRNA modifications, we examined fragment type, length, boundary position, and coverage profiles across age groups. We first assessed correlations between fragment abundance and age for individual tRNA isodecoders, stratified by fragment type (Fig. 2A; GSE111279). Among the top 15 most abundant nuclear-encoded tRFs, positive Spearman correlations predominated across all three fragment classes (5′, internal, and 3′ tRFs), with several reaching statistical significance (p < 0.05). In contrast, mitochondrial-encoded tRFs largely displayed negative or near-zero correlations with age, indicating that the age-associated tRF increase is specific to nuclear-encoded tRNAs. Similar concordant increases in nuclear tRFs with age were replicated in the independent datasets. However, the decrease in mt-tRFs was not, with both datasets instead showing no significant change (Sup. Fig. 3A).

**Figure 2.**
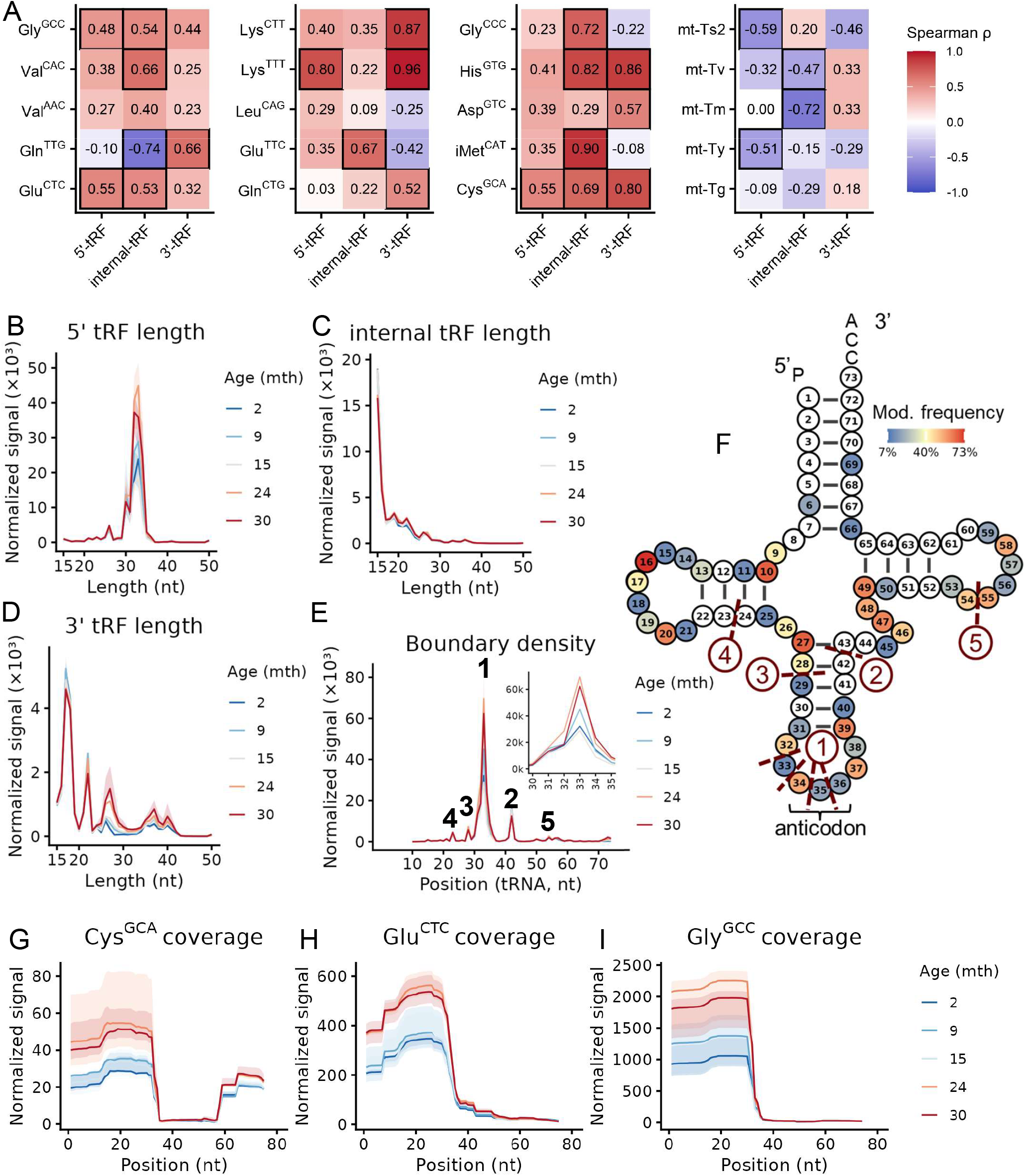
Age-associated elevation of tRFs is consistent with increased cleavage at specific sites. **(A)** Spearman correlation (ρ) between fragment abundance and age for individual tRNA isodecoders in mouse brain (GSE111279), stratified by fragment type (5′, internal, and 3′ tRFs). The top 15 most abundant nuclear-encoded and top 5 most abundant mitochondrial-encoded tRFs are shown. Values represent per-fragment correlations across ages. Solid black borders indicate FDR-adjusted p < 0.05 (Benjamini-Hochberg). **(B-D)** Length distributions of (B) 5′, (C) internal , and (D) 3′ tRFs across age groups, shown as DESeq2-normalized signal per fragment length. Lines represent median signal across fragments of each length; shaded regions indicate IQR. **(E)**Fragment boundary density across the tRNA body, plotted as DESeq2-normalized signal versus pseudo-Sprinzl position (nt; 5′ to 3′, positions 1-76). Each position reflects the summed signal of fragments whose 3′ end maps to that position. Lines represent median signal; shaded regions indicate IQR. A cluster of peaks at positions 32-38 nt shows a progressive increase in signal with age. Four additional peaks are indicated. **(F)**Canonical tRNA cloverleaf secondary structure annotated with pseudo-Sprinzl positions (1-76 nt) and modification frequencies from MODOMICS. Cleavage sites enriched in panel E (red marks; numbered) are mapped onto defined structural regions. The dominant cluster at positions 32-38 nt corresponds to the anticodon loop. **(G-I)** Nucleotide-resolution coverage maps for representative nuclear-encoded tRNAs: (G) Cys^GCA^, (H) Glu^CTC^, and (I) Gly^GCC^ (5′ to 3′). Lines represent median DESeq2-normalized read coverage across positions; shaded regions indicate IQR.

Analysis of fragment length distributions revealed distinct size profiles for each tRF class (Fig. 2B-D). 5′ tRFs were predominantly enriched in the 30-40 nt range, with signal in this size class increasing progressively with age (Fig. 2B). Internal tRFs displayed a markedly different profile, with the majority of fragments clustering at ∼15 nt, near the lower size cutoff applied during filtering (Fig. 2C). Notably, some internal tRFs may represent incomplete reverse transcription products rather than genuine cleavage products. For example, short 15-20 nt internal tRFs may originate from the angiogenin cleavage site, with reverse transcriptase drop-off occurring at the heavily modified ∼16 nt position (Fig. 2F, mouse tRNA modification data obtained from Modomics^25^), though this remains to be tested. 3′ tRFs exhibited a broader size distribution, with multiple enriched size classes spanning 15-45 nt, potentially reflecting heterogeneous cleavage or modification positions along the 3′ portion of the tRNA (Fig. 2D). Across all three fragment types, dominant size classes were preserved across age groups with increased signal intensity but stable peak positions, arguing against non-specific RNA degradation as the primary driver of age-associated tRF accumulation. Consistent with this interpretation, fragment boundary mapping confirmed that cleavage sites are maintained with age (Fig. 2E-F). Of the five major boundary peaks, the dominant peak corresponds to the anticodon loop (32-38 nt), the canonical angiogenin cleavage site^24^, and shows an age-dependent increase. The remaining peaks are comparatively smaller. Similar patterns were observed in an independent dataset (Sup. Fig. 3B-E).

Coverage maps for representative nuclear-encoded tRNAs, Cys^GCA^, Glu^CTC^, and Gly^GCC^, further supported this conclusion (Fig. 2G-I). Coverage profiles were qualitatively similar across age groups, with elevation in intensity with age. The most pronounced increase occurred between 15 and 24 months in mice, a period during which C57BL/6 mice begin to exhibit considerable cognitive impairment^26^, and which corresponds to the transition from middle to old age in humans^27^. Higher signal at the 5′ fragment region and a sharp boundary at the anticodon loop (32∼38 nt) were consistent with angiogenin-mediated cleavage.

To explore why some nuclear tRF families are not age-responsive, we compared fragment boundary profiles of the top 10 age-elevated versus top 10 age-stable families, aggregated across 17 brain and brain region samples from all three datasets (Sup. Fig. 3F). Age-elevated families consistently showed higher boundary signal at the angiogenin cleavage site than age-stable families (Sup. Fig. 3G-I), suggesting that susceptibility to angiogenin cleavage is a key determinant of age-associated tRF accumulation. In contrast, fragment boundary profiles for mitochondrial tRNAs (Supp. Fig. 3J-K) showed strong signal at the angiogenin cleavage site in both brain datasets, particularly GSE111279, but with age-dependent decrease or no change. As mitochondrial tRNAs are largely confined within the mitochondrial matrix, their inaccessibility to cytoplasmic angiogenin may account for this lack of age-dependent increase. Together, these data demonstrate that age-associated tRF accumulation reflects amplified cleavage at conserved sites, with angiogenin as a major contributor, rather than nonspecific tRNA degradation.

### Validation of tRF upregulation in the aging mouse brain

Quantification of tRFs from small RNA sequencing data presents several technical challenges. Most small RNA-seq protocols are optimized for miRNAs of ∼22 nt with ligatable 5’-P and 3’-OH ends, including those used to generate the public datasets reanalyzed here (Sup. Table S1). However, tRFs are typically ∼30-38 nt in length, have different end modifications and more extensive internal modifications (ref. Fig. 2F), therefore introducing biases in sequencing-based quantification^28^. Therefore, independent validation of RNA-seq-based tRF quantification using orthogonal approaches is essential. We first isolated RNA from multiple mouse organs and performed SYBR Gold staining of total RNA on urea-polyacrylamide gels, which revealed lower intensity of RNA bands in the ∼25-50 nt range in brain regions relative to other organs (Sup. Fig. 4A). Subsequent northern blotting with a 5′Cys^GCA^ probe confirmed a dominant 5′ tRF band at ∼30 nt, with brain regions exhibiting the lowest signal compared to liver, kidney, heart, spleen, and lung (Sup. Fig. 4B).

To validate age-associated tRF upregulation, we analyzed RNA isolated from the whole brain of young (5 months) and old (24+ months) male mice using a series of complementary quantification approaches. SYBR Gold staining of total RNA revealed three distinct size classes corresponding to full-length tRNAs (FL tRNAs; 70-80 nt), tRFs (30-40 nt), and miRNAs/other small RNAs (17-25 nt) (Fig. 3A). Notably, the distribution of bands elevated in aged brain tissue corresponded to those observed in cells treated with exogenous angiogenin (Fig. 3A). Quantification confirmed that FL tRNA and 17-25 nt miRNA/small RNAs were unchanged, while tRFs in the 30-40 nt range were significantly elevated in aged brains (Fig. 3B-D). These findings are consistent with increased angiogenin-mediated tRNA cleavage in the aging brain. Northern blotting using tRF-specific probes confirmed that 5′Cys^GCA^, 5′Glu^CTC^, and 3′Glu^CTC^ were all significantly upregulated in aged relative to young brains (Fig. 3E-G), while their corresponding parental full-length tRNAs remained unchanged. Neither FL tRNAs nor tRFs derived from Glu^CTC^ and Cys^GCA^ were changed in young versus old spleen (Sup. Fig 4C-E). As 5′ and 3’Gly^GCC^ could not be detected by Northern blot, we turned to the tRF-specific RT-qPCR for further analysis. When normalized to the housekeeping miR-99a, RT-qPCR results were consistent with northern blot findings, validating the approximately two-fold upregulation of 5′Cys^GCA^ and 5′Glu^CTC^ in aged brains. Using the same approach, we further detected significant increases in 5′Glu^TTC^, 5′Gly^CCC^, and 5′Gly^GCC^ (Fig. 3H-L), providing additional support for the global increase in tRF abundance observed in the small RNA-seq dataset.

**Figure 3.**
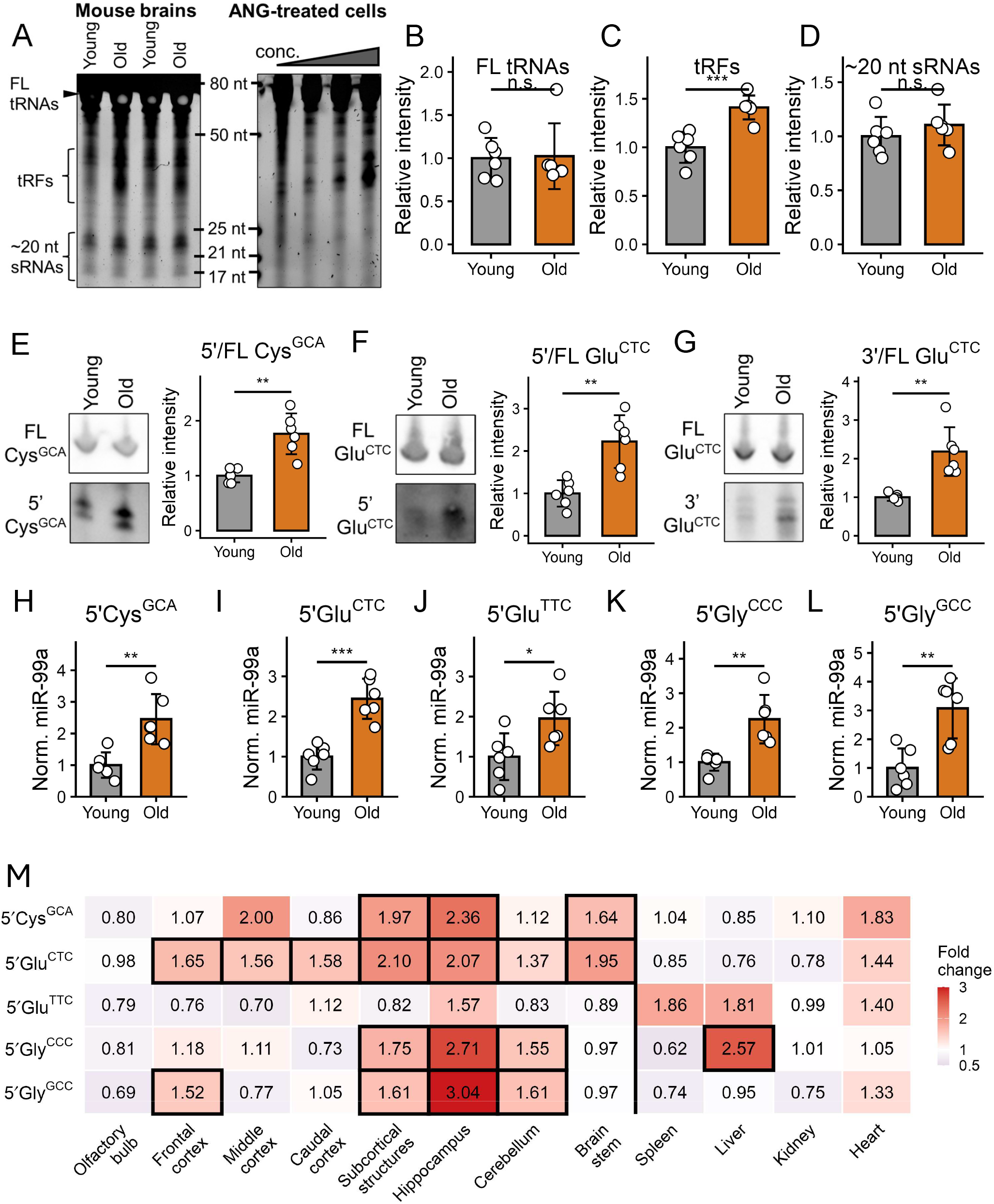
Validation of tRF upregulation in the aging brain. **(A)** Left: SYBR Gold staining of 10 µg total RNA isolated from brains of young (5 months) and old (24+ months) mice. Right: SYBR Gold staining of 3 µg total RNA from SK-N-AS cells treated with 0, 0.1, 0.25, and 0.5 µg/mL exogenous angiogenin for 1 hour. **(B-D)** Quantification of signal within defined size windows from the SYBR Gold gel: (B) full-length tRNAs (FL; 70-80 nt), (C) tRNA fragments (tRFs; 25-50 nt), and (D) ∼20nt sRNAs (17-25 nt). **(E-G)** Representative northern blot images and quantification of tRF band intensity relative to the corresponding full-length tRNA band. **(H-L)** RT-qPCR quantification of individual tRFs, normalized to housekeeping miR-99a. **(M)** RT-qPCR quantification of individual tRFs across brain regions and peripheral organs, normalized to miR-99a. Values represent fold change relative to young controls. Black borders indicate statistically significant differences. For panels B-M, statistical significance was determined using an unpaired two-tailed t-test (n = 6 male mice per group). *p < 0.05; **p < 0.01; ***p < 0.001. Error bars represent SD.

Finally, to assess whether age-associated tRF upregulation was restricted to particular brain regions or broadly distributed across the brain, we profiled five tRF species across 8 brain regions and 4 peripheral organs by RT-qPCR (Fig. 3M, source data provided). Across all brain regions examined, tRF levels were broadly elevated in aged mice relative to young controls. Notably, 5′Glu^CTC^ was significantly upregulated in 7 of 8 brain regions. The hippocampus and subcortical structures showed significant increases in 4 of 5 tRFs examined, while the olfactory bulb showed no significant changes. Among peripheral tissues, 5′Gly^CCC^ was significantly elevated in the liver, a trend toward increased 5’Cys^GCA^, 5′Glu^CTC^, and 5′Gly^GCC^ was observed in the heart (p = 0.07-0.1), and no significant age-associated changes were detected in the kidney or spleen. Taken together, these data indicate that a global age-associated upregulation of tRFs is a brain-specific phenomenon, broadly consistent with findings from small RNA-seq datasets (Fig.1).

### tRFs are Upregulated in Chronic and Acute Human Brain Diseases

As small RNA-seq data from healthy aging human brain are not readily available, we instead examined whether the tRF upregulation observed in mouse brain aging is recapitulated in human neurological diseases. We analyzed publicly available frontal lobe small RNA-seq data (E-MTAB-12731) from donors with frontotemporal dementia (FTD) subtypes (FTD-C9orf72, FTD-GRN, and FTD-MAPT) compared to neurologically normal controls (n = 8-13 donors per group). Differential expression analysis revealed that tRF subclasses exhibited a strikingly high fraction of significantly upregulated members across all three subtypes, consistently exceeding upregulation rates for miRNAs as reference (Fig. 4A, Sup. Fig. 5A). This effect was most pronounced in FTD-MAPT, where ∼75% of 5’ tRFs were significantly upregulated, but was also observed across the other two subtypes, suggesting a shared disease-associated response rather than a mutation-specific effect. 5’ tRF length distributions showed a dominant peak at ∼33 nt across all groups, with substantially increased signal in FTD donors relative to controls (Fig. 4B). Fragment boundary density analysis confirmed a sharp cleavage peak at the expected angiogenin cleavage site in the anticodon loop (Fig. 4C). Coverage maps for three representative nuclear-encoded tRNAs, Cys^GCA^, Glu^CTC^ and Gly^GCC^, were consistent with selective 5’ end enrichment and showed elevated signal in FTD groups relative to controls, again strongest in FTD-MAPT (Fig. 4D-F). Northern blot analysis of human cortex samples from an independent cohort (Sup. Table S2) confirmed the upregulation of 5′Glu^CTC^, and 5′Gly^GCC^, revealing two discrete ∼30-35 nt bands at significantly higher intensity in patients with frontotemporal lobar degeneration with tau pathology (FTLD-tau) (Fig. 4F-G; all samples and quantification in Sup. Fig. 5D-L). Similar to mouse brain aging, boundary density analyses suggest that tRF families unchanged in FTD cortexes are less susceptible to angiogenin cleavage than those that are upregulated, while mitochondrial tRNAs, despite high signal at the angiogenin cleavage site, show no differences across groups (Sup. Fig. 5A-C).

**Figure 4.**
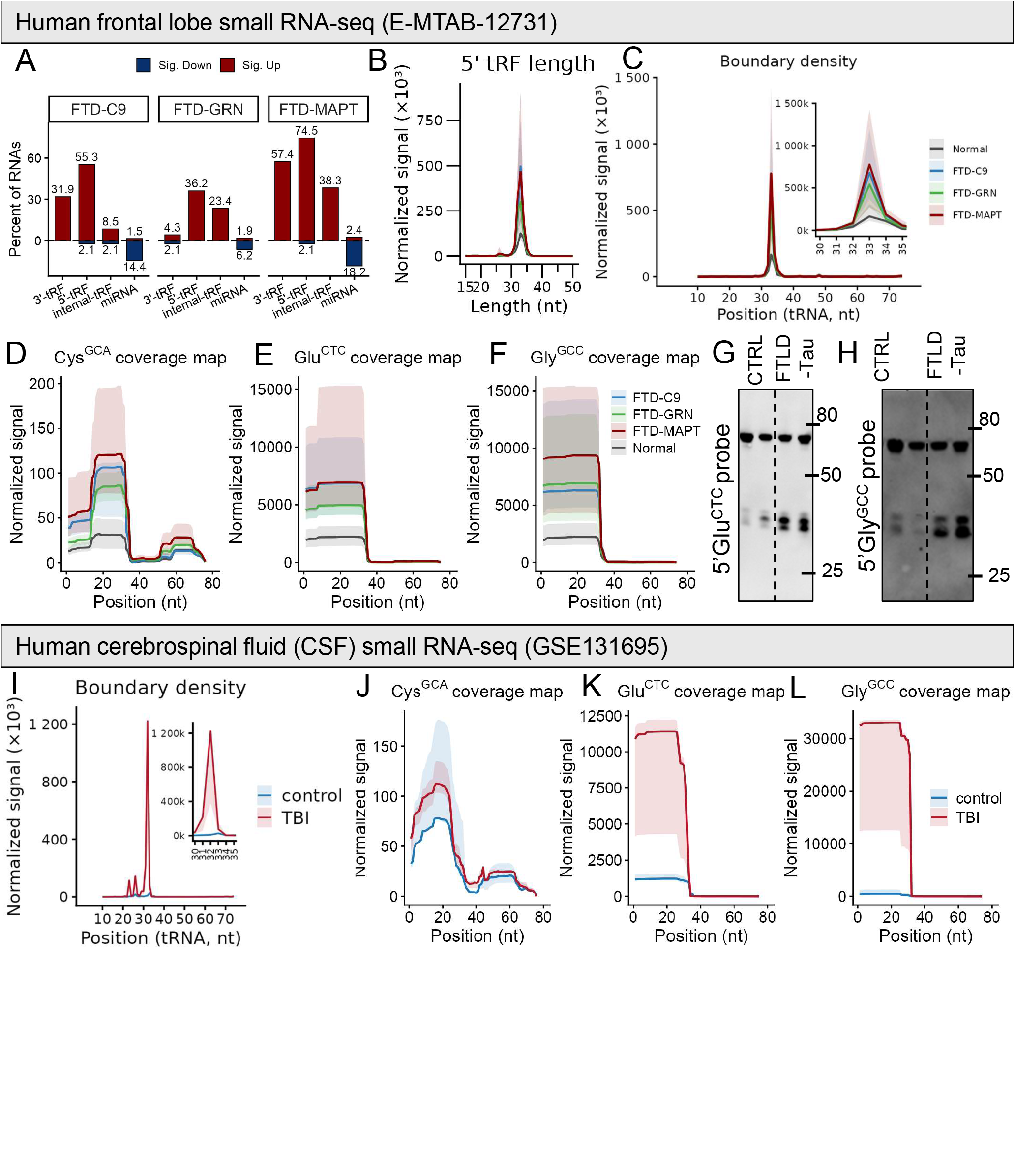
tRFs are upregulated in chronic and acute brain disease. **(A-F)** Human frontal lobe small RNA-seq data (E-MTAB-12731, batch 1; n = 8-13 donors per group). **(A)**Fraction of RNAs within each biotype with significant upregulation (red) or downregulation (blue) in each FTD subtype (C9orf72, GRN, MAPT) relative to normal controls. Significance was defined by DESeq2 (padj < 0.05,|log2FC| > 0.5). **(B)**Length distribution of 5′ tRFs shown as DESeq2-normalized signal per fragment length across FTD subtypes and normal controls. Lines represent median signal; shaded regions indicate IQR. A dominant peak at ∼33 nt is present across all groups. **(C)**Fragment boundary density across the tRNA body (5′ to 3′), plotted as DESeq2-normalized signal versus pseudo-Sprinzl position. Lines represent median signal; shaded regions indicate IQR. A consistent peak at positions 32-38 nt, corresponding to the anticodon loop, is observed across all groups. **(D-F)** Nucleotide-resolution coverage maps for representative nuclear-encoded tRNAs (D) Cys^GCA^, (E) Glu^CTC^ and (F) Gly^GCC^ (5′ to 3′). Lines represent median DESeq2-normalized read coverage across positions; shaded regions indicate IQR. **(G-H)** Representative Northern blot images of RNA isolated from human cortex samples and probed for (G) 5′Glu^CTC^ and (H) 5′Gly^GCC^, showing bands at ∼30-35 nt across normal controls and FTLD-Tau samples. **(I-L)** Human cerebrospinal fluid (CSF) small RNA-seq data (GSE131695; n = 5 per group, control versus 24-hour post-TBI). **(I)** Fragment boundary density across the tRNA body, plotted as DESeq2-normalized signal versus pseudo-Sprinzl position. Lines represent median signal; shaded regions indicate IQR. A peak at positions 32-38 nt, corresponding to the anticodon loop, is significantly elevated in the TBI group. **(J-L)** Nucleotide-resolution coverage maps for (J) Cys^GCA^,(K) Glu^CTC^ and (L) Gly^GCC^. Lines represent median DESeq2-normalized read coverage across positions; shaded regions indicate IQR. For all panels, median DESeq2-normalized counts are shown; shading indicates IQR.

To determine whether tRF upregulation extends beyond chronic neurodegeneration to acute brain injury, we analyzed cerebrospinal fluid (CSF) small RNA-seq data (GSE131695) from patients sampled 24 hours after traumatic brain injury (TBI) versus controls (n = 5 per group). Fragment boundary density in TBI CSF showed a cleavage peak at the same tRNA body position as observed in frontal lobe tissue, with approximately 1000-fold increased signal relative to controls (Fig. 4I). While Cys^GCA^ showed no change, coverage maps Glu^CTC^ and Gly^GCC^ confirmed markedly elevated 5′ tRF signal in TBI samples, with the characteristic drop-off at the cleavage site, (Fig. 4J-L). Finally, we identified nuclear tRNA families that show no changes with aging or neurological disease, represented here by Leu^TAG^, Leu^CAA^, Ala^AGC^, Tyr^GTA^, and Ser^GCT^, in contrast to the age- and disease-responsive families Glu^CTC^, Gly^GCC^, Gly^CCC^, Lys^TTT^, and Cys^GCA^. Age-stable families carried one or more modifications within the anticodon loop, whereas most age-responsive families had none (Sup. Fig. 5M, human tRNA modification data obtained from MODOMICS^25^), consistent with the hypothesis that anticodon loop modifications reduce accessibility to angiogenin cleavage and thereby limit fragment production.

## DISCUSSION

Here we report that nuclear-encoded tRNA-derived fragments accumulate selectively in the aging mouse brain. This finding was replicated across three independent small RNA-seq datasets and validated by orthogonal methods including northern blotting and RT-qPCR. The age-associated increase was specific to nuclear-encoded tRFs, broadly distributed across brain regions, and largely absent from peripheral organs. Fragment length distributions, cleavage boundary profiles, and coverage maps of nuclear tRFs were all consistent with cleavage at conserved, structurally defined sites rather than nonspecific RNA degradation. Furthermore, analogous tRF upregulation was observed in human brain tissue from FTD donors and in CSF from TBI patients, suggesting that the tRF response seen during normal aging is further amplified under conditions of acute and chronic neurological stress. Most datasets included samples from both sexes but were underpowered to detect sex-specific effects. Experimental validation was performed in male mice and in human tissue from donors of both sexes. While tRF upregulation in the brain is likely to occur across both sexes, any variation in magnitude and impact remains to be investigated.

Several technical caveats warrant consideration. Standard small RNA-seq library preparation protocols are optimized for miRNA-sized species (∼22 nt) and may underrepresent tRFs, which are typically 30-35 nt. More importantly, tRNAs carry extensive base modifications with an estimated 43 distinct chemical modifications and an average of 13 modifications per tRNA molecule^8^, which can impede reverse transcription and introduce systematic biases in sequencing-based quantification^28^. For example, “hard-stop” modifications such as m1A, m1G, m22G and m3C can block reverse transcriptase, leading to premature termination of sequencing reads. Indeed, we hypothesize that some fragments classified as internal tRFs, or fragments < 15nt filtered out during trimming, may represent biological 5′ or 3′ tRFs prematurely truncated during library construction. While the general trends reported here are robust, abundance estimates for specific tRF species should be interpreted with caution, and validation by orthogonal methods remains important. In addition, while we collapse isodecoders into family-level features to reduce redundancy from multimapping reads, this may obscure functional differences between closely related species differing by as little as a single nucleotide. Continued development of modification-aware sequencing^28^, such as ARM-seq^29^ or PANDORA-seq^30^ alongside isodecoder-level quantification methods will be essential for a more complete picture of the tRF landscape.

The fragment boundary and coverage data presented here strongly implicate angiogenin as a primary driver of tRF accumulation. Angiogenin is a stress-activated ribonuclease that cleaves tRNAs at the anticodon loop to generate 5’ and 3’ tRNA halves^5,24^. Loss-of-function mutations in angiogenin, and rarely gain-of-function variants, have been linked to decreased motor neuron survival and amyotrophic lateral sclerosis^31,32^. The dominant cleavage peak across all datasets is consistent with angiogenin activity. Age-stable families also carry anticodon loop modifications absent in age-responsive families, potentially resulting in lower boundary density at this site and reduced susceptibility to cleavage. Conversely, mitochondrial tRFs showed negative or near-zero age and disease correlations despite high boundary density at the anticodon loop, consistent with their retention within the mitochondrial matrix rendering them largely inaccessible to cytoplasmic nucleases. However, the additional boundary peaks identified suggest that other RNases may also contribute, potentially including members of the RNase A superfamily or other tRNA endonucleases. The age-associated increase in tRF levels could reflect increased angiogenin expression, relocalization of angiogenin from the nucleus to the cytoplasm^15,33^, or reduced activity of angiogenin inhibitors^34^ or tRF-degrading nucleases. Decreased tRNA modification levels, as reported in the cortex of 5XFAD mice^35^, may further increase substrate accessibility and promote cleavage. What remains to be explained is why this response appears to be brain-specific. The brain may be particularly susceptible due to its high metabolic demands, limited regenerative capacity, and progressive accumulation of cellular stress with age. Whether angiogenin activity or localization, or tRNA susceptibility to cleavage change specifically in brain tissue with age, and what upstream signals, beyond age-associated stress, drive this, are important open questions.

tRNAs are among the most evolutionarily conserved molecules in biology, and this conservation extends to the cleavage site positions and fragment size profiles observed across mouse and human samples. Nevertheless, absolute tRF abundance may matter functionally, and we observe an interesting quantitative discrepancy exists between mice and humans. Among the various small RNA-seq databases included in this study, tRF reads constitute approximately 2% of small RNA-seq reads in mouse brain, compared to roughly 10-25% in human datasets (Supp. Fig. 1B, 2A, and 5A). Consistently, northern blotting required substantially less total RNA from human samples (0.5 µg) than from mouse brain (10 µg) to detect specific tRFs. It is difficult to disentangle how much of this difference reflects genuine species-level biology versus technical factors such as postmortem interval, RNA quality, and tissue handling. Notably, a recent small RNA-seq study comparing tissues across species reported ∼2% tRNA-derived reads in mouse brain and ∼20% in rhesus macaque brain^36^, raising the possibility that tRF abundance is genuinely elevated in primate brain independent of sample quality. Whether this apparent increase reflects differences in cleavage activity, tRF stability, or other species-specific factors warrants further investigation.

Perhaps the most pressing question raised by these findings is whether accumulating tRFs are functionally inert byproducts of stress-induced cleavage or active contributors to brain aging and neurological disease. Evidence from our own work and other recent studies suggest the latter. We have reported that 5’Glu^CTC^ and 5’Gly^GCC^ are highly elevated in AD and FTD brains, and in corresponding cell and animal models^20^. These tRF species can promote tau aggregation and fibrilization both in biophysical assays and in cultured neurons, directly linking them to pathological processes relevant to neurodegeneration^20^. Beyond tau aggregation, 5’Glu^CTC^ has been reported to impair mitochondrial functions, leading to age-related memory decline in mice^15^. Other tRFs have also been reported to modulate translation, interact with RNA-binding proteins, and regulate gene expression through miRNA-like mechanisms^7^, any of which could have downstream consequences in the aging brain. Determining which tRF species are functionally active, how they contribute to brain pathology, and whether they can be targeted therapeutically will be of particular interest going forward.

Overall, we propose that tRF accumulation represents a conserved feature of brain aging and select neurological diseases, with upstream regulatory mechanisms, downstream functional consequences, and condition-specific tRF signatures that remain to be further investigated (Fig. 5) Future work should prioritize several directions. Identifying the specific nucleases responsible for tRF generation in the aging brain and determining what drives their increased activity or altered localization, will be critical for understanding upstream mechanisms. Systematic functional characterization of the age-associated tRF species is needed to establish which tRFs contribute causally to neuronal dysfunction or neurodegeneration. Given the elevation of tRFs in both FTD and TBI, it will also be important to assess whether tRF levels in CSF or other biofluids could serve as accessible biomarkers of brain stress or injury. Notably, the therapeutic potential of targeting tRFs is already supported by experimental evidence. We have shown that ASO-mediated knockdown of 5’Glu^CTC^ can reduce tau aggregation and improve neuronal health^20^, and independent work has demonstrated that targeting 5’Glu^CTC^ decreases age-related cognitive impairment in mice^15^. Together, these findings suggest that reducing tRF accumulation may be a tractable strategy for mitigating brain aging and neurological disease.

**Figure 5.**
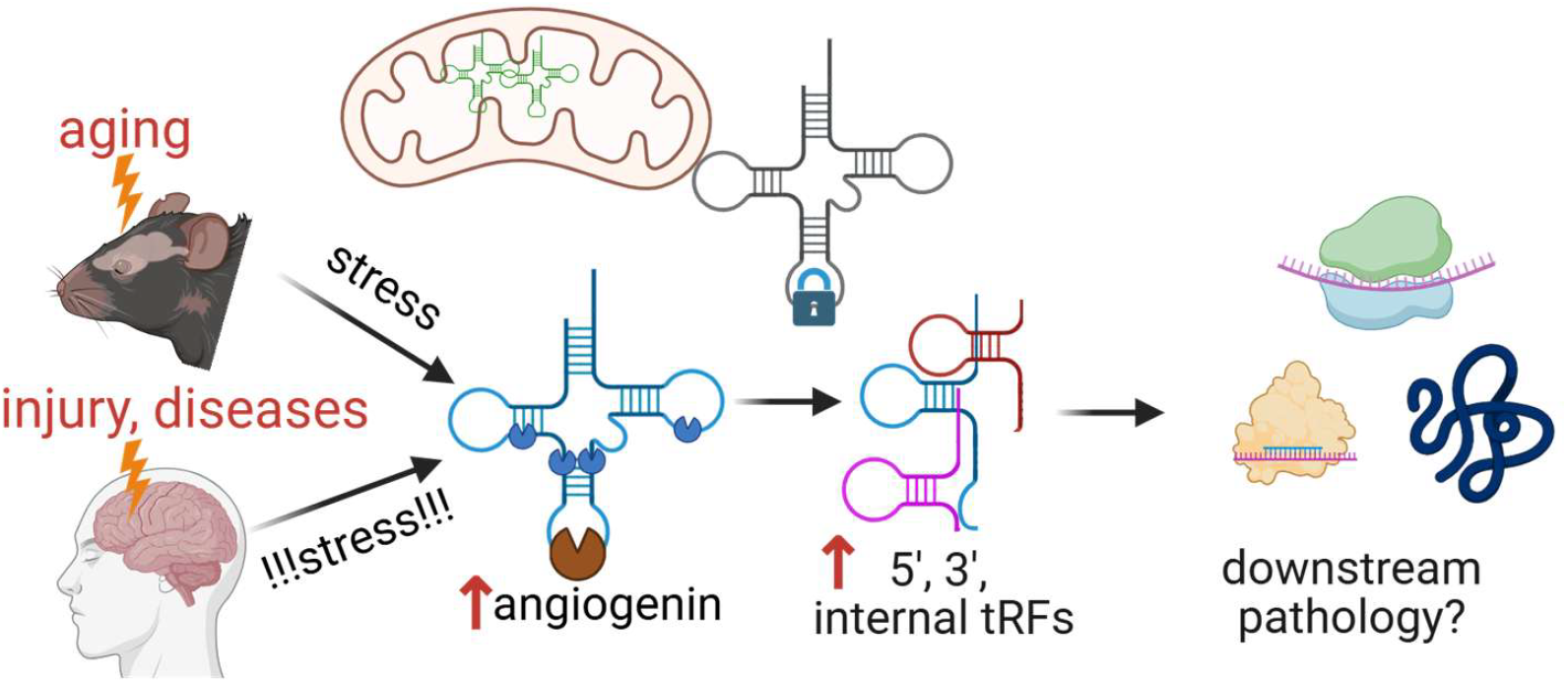
Proposed model for tRF accumulation in the aging and diseased brain. Aging is associated with selective tRF accumulation in the brain. Acute or chronic brain diseases may represent an amplified form of this response. Stress-induced nuclease activity, particularly by angiogenin, promotes cleavage of nuclear-encoded tRNAs and accumulation of 5′, 3′, and internal tRFs. Mitochondrial tRNAs, which are retained within the mitochondrial matrix, and nuclear-encoded tRNAs with lower cleavage site accessibility remain relatively stable across age and disease. Which specific tRFs contribute to downstream pathology, and through what mechanisms, including impaired translation, protein aggregation, or miRNA-like gene silencing, remain to be determined. Figure was created with biorender.com.

## Funding sources

This work was supported by the National Institute on Aging (RF1AG090598 to A.M.K.) and the Rainwater Charitable Foundation (A.M.K.). A.K was supported by Uehara Memorial Foundation and the Japan Society for the Promotion of Science.

## Author contribution

L.D.N. conceived and designed the study, performed experiments, conducted computational analyses, and wrote the manuscript. A.K. contributed to early conception and performed preliminary experiments. A.N. curated and assisted with human sample characterization. V.K. facilitated human brain tissue procurement and provided reagents. P.I. provided expertise, reagents, and contributed to manuscript editing. A.M.K. conceived and supervised the study, acquired funding, and critically reviewed and edited the manuscript. All authors reviewed and approved the manuscript.

## Acknowledgement

We thank the National Institute on Aging (NIA) Aged Rodent Colonies (Division of Aging Biology, Translational Research Branch) for providing aged mice. We thank the staff at the Massachusetts General Brigham (MGB) Brain Bank for facilitating the acquisition of human brain tissue. We thank Drs. Erik J. Uhlmann (BIDMC), Evgeny Deforzh (BWH), Yanhong Zhang, and Zelong Zheng (BWH) for helpful discussions and assistance with experiments. Figure 5 was created with BioRender.com.

## Conflict of interest

All authors declare no competing interests.

## Data and Code Availability

Raw read count tables and analysis code for reanalyzed datasets are deposited on Zenodo (DOI: 10.5281/zenodo.20141089). The small RNA-seq alignment pipeline is available on GitHub (https://github.com/Krichevsky-Lab/smallRNA_seq_pipeline; DOI: 10.5281/zenodo.20140745). Original sequencing data are available under accession numbers listed in Supp. Table S1. Source data for Fig. 3 and Sup. Figs. 4 and 5, along with full northern blot and SYBR Gold gel images, are provided as supplementary materials.

## Supplementary Figure Legend

**Supplementary Figure 1.**
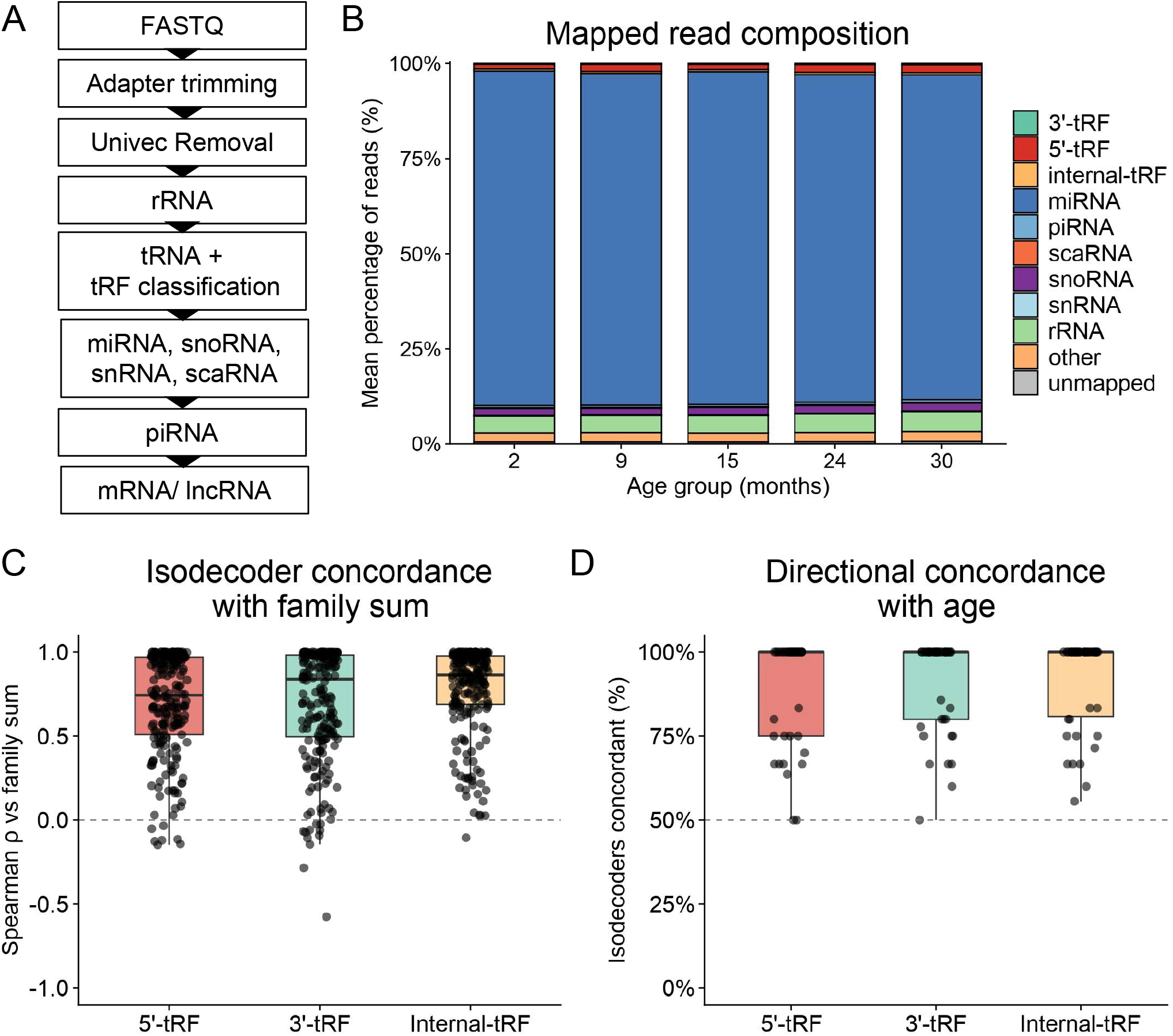
Small RNA-seq alignment pipeline and tRF isodecoder concordance. **(A)** Schematic of the sequential alignment strategy used for small RNA-seq read processing. **(B)** Distribution of percentage mapped reads by RNA biotype across age groups, showing overall mapping efficiency. **(C)** Spearman correlation of each tRF isodecoder with the summed expression profile of its corresponding tRF family across samples. Lines represent median ρ; shading indicates IQR. **(D)** Directional concordance of age association for isodecoders within the same tRF family, shown as the fraction of isodecoders sharing the same direction of Spearman correlation with age as their family sum.

**Supplementary Figure 2.**
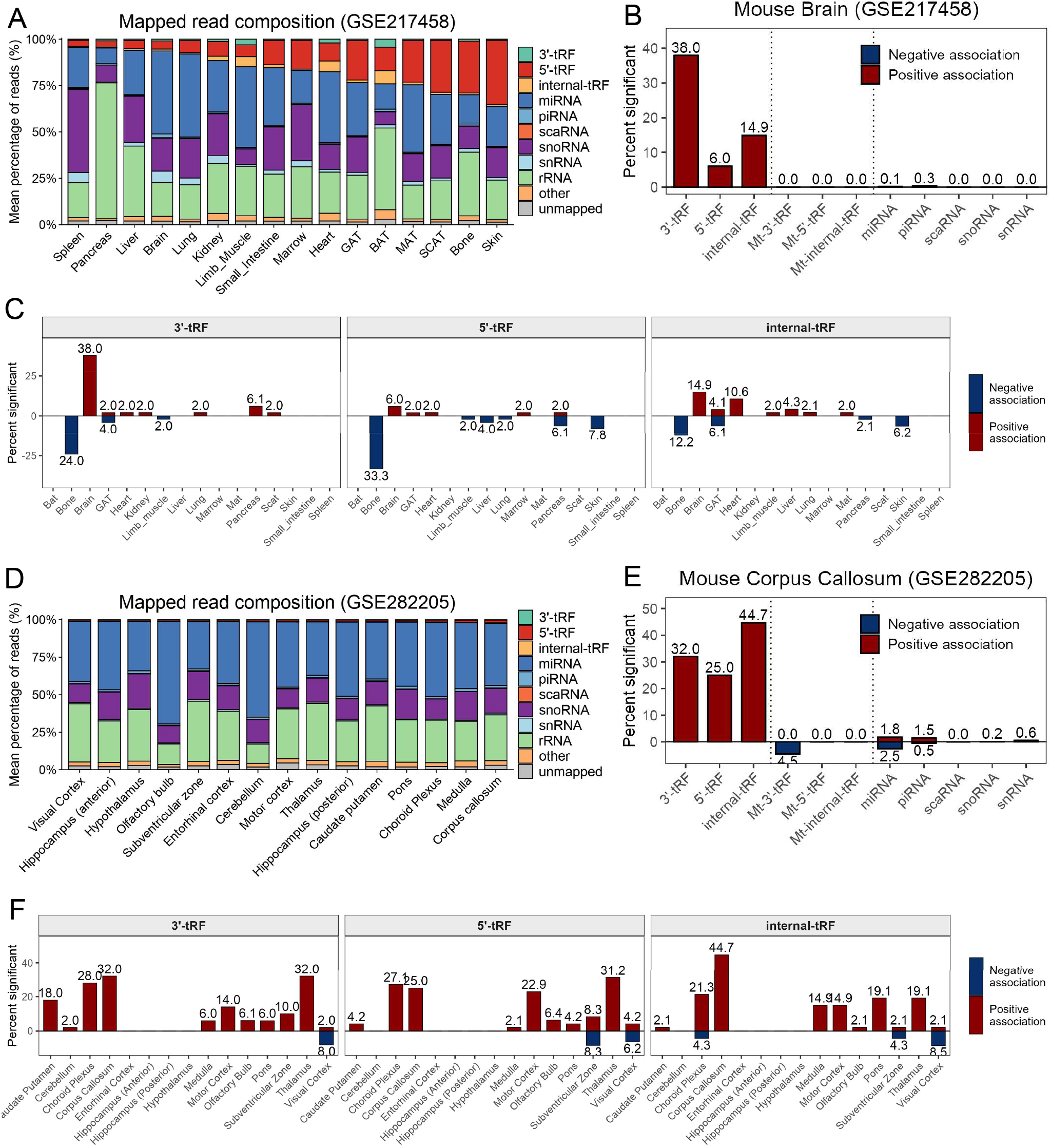
Organ and brain region specificity of age-associated tRF accumulation. **(A)** Mean percentage of reads mapped to each RNA biotype across organs (GSE217458; 10 age groups, n = 4-6 male and female mice per group), ordered from lowest to highest total tRF content. **(B)** Percentage of RNAs within each biotype with a significant positive (red) or negative (blue) Spearman correlation with age in mouse brain (GSE217458). Significance threshold: FDR-adjusted p < 0.05 (Benjamini-Hochberg). **(C)** Percentage of RNAs within each biotype showing significant positive (red) or negative (blue) age association, stratified by organ and tRF class (3′, 5′, and internal tRFs). **(D)** Mean percentage of reads mapped to each RNA biotype across brain regions (GSE282205; 8 age groups, n = 6-10 male and female mice per group), ordered from lowest to highest total tRF content. **(E)** Percentage of RNAs within each biotype with a significant positive (red) or negative (blue) Spearman correlation with age in the corpus callosum (GSE282205). Significance threshold: FDR-adjusted p < 0.05 (Benjamini-Hochberg). **(F)** Percentage of RNAs within each biotype showing significant positive (red) or negative (blue) age association, stratified by brain region and tRF class (3′, 5′, and internal tRFs).

**Supplementary Figure 3.**
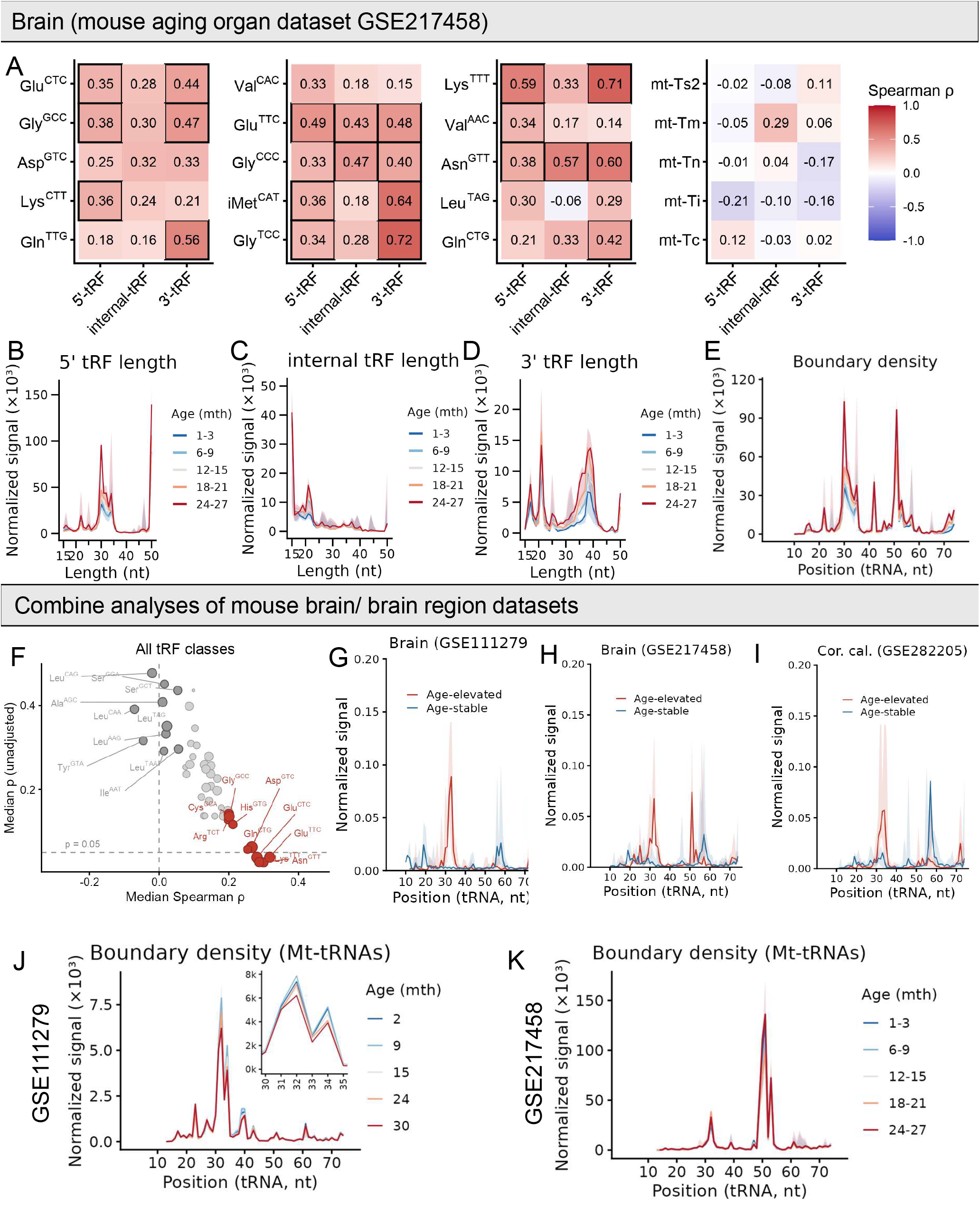
Age-associated tRF accumulation in brain reflects increased cleavage at specific sites. **(A)** Spearman correlation (ρ) between fragment abundance and age for individual tRNA isodecoders in mouse brain (GSE217458), stratified by fragment type (5′, internal, and 3′ tRFs). The top 15 most abundant nuclear-encoded and top 5 most abundant mitochondrial-encoded tRFs are shown. Solid black borders indicate FDR-adjusted p < 0.05. Values represent per-fragment correlations across ages. **(B-D)** Length distributions of (B) 5′, (C) internal, and (D) 3′ tRFs across age groups (GSE217458), shown as DESeq2-normalized signal per fragment length. Lines represent median signal; shaded regions indicate IQR. **(E)**Fragment boundary density across the nuclear-encoded tRNA body, plotted as DESeq2-normalized signal versus pseudo-Sprinzl position (nt; 5′ to 3′). Lines represent median signal; shaded regions indicate IQR. A cluster of peaks at positions 32-38 nt, corresponding to the anticodon loop, shows a progressive increase in signal with age. Notably, a substantial fraction of 5′-tRFs reaches the maximum library read length of 50 nt (panel B), corresponding to a boundary density peak at ∼50 nt (pseudo-Sprinzl). These fragments may represent full-length tRNAs co-sequenced with tRFs. **(F)**Median Spearman ρ versus median unadjusted p-value per tRNA family (all tRF classes combined), aggregated across 17 mouse brain and brain region samples from three datasets. The top 10 families with the highest median ρ (age-elevated, red) and the 10 families with median ρ closest to 0 (age-stable, grey) are labeled. **(G-I)** Fragment boundary density for the top 10 age-elevated versus top 10 age-stable tRNA families in (G) brain hemisphere (GSE111279), (H) brain hemisphere (GSE217458), and (I) corpus callosum (GSE282205). Boundary density is normalized per tRNA family before averaging. Lines represent median signal; shaded regions indicate IQR. Age-elevated families show higher boundary density at positions 32-38 nt than age-stable families across all three datasets. **(J-K)** Fragment boundary density across the mitochondrial-encoded tRNA body, plotted as DESeq2-normalized signal versus pseudo-Sprinzl position (nt; 5′ to 3′). Lines represent median signal; shaded regions indicate IQR. A cluster of peaks between approximately 32-38 nt shows a progressive decrease or no change with age.

**Supplementary Figure 4.**
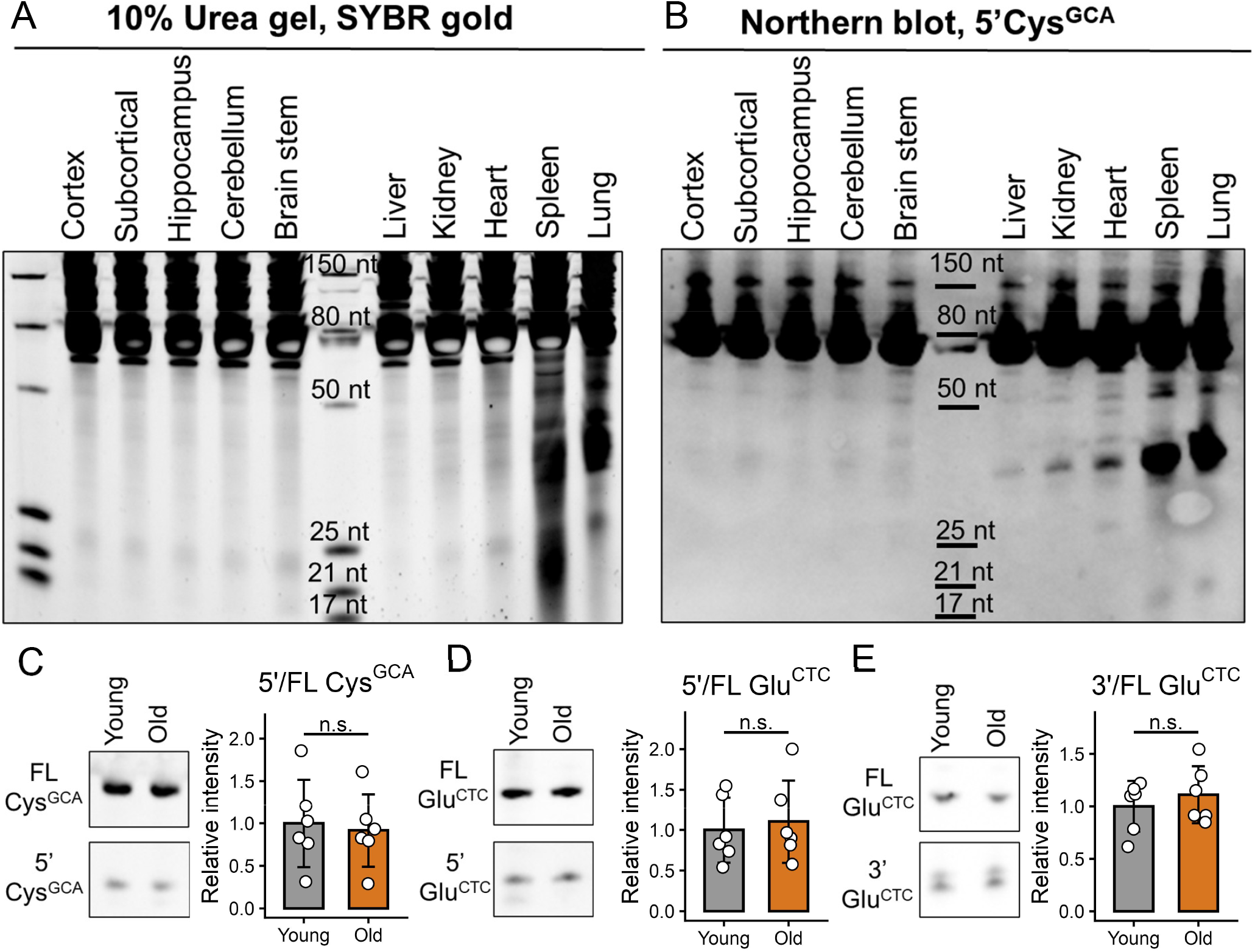
Quantification of tRF expression across organs. **(A)** Representative SYBR Gold staining of 5 µg total RNA isolated from different brain regions and solid organs of a young (5-month-old) male mouse. **(B)** Northern blot hybridization for 5′Cys^GCA^ performed on the same samples as in panel A. **(C-E)** Representative northern blot images and quantification of tRF band intensity relative to the corresponding full-length tRNA band in spleen. Each lane contains 0.5 µg total RNA isolated from young (5 months) and old (≥24 months) mice. Statistical significance was determined using an unpaired two-tailed t-test (n = 6 mice per group). *p < 0.05; **p < 0.01; ***p < 0.001; ****p < 0.0001. Error bars represent SD.

**Supplementary Figure 5.**
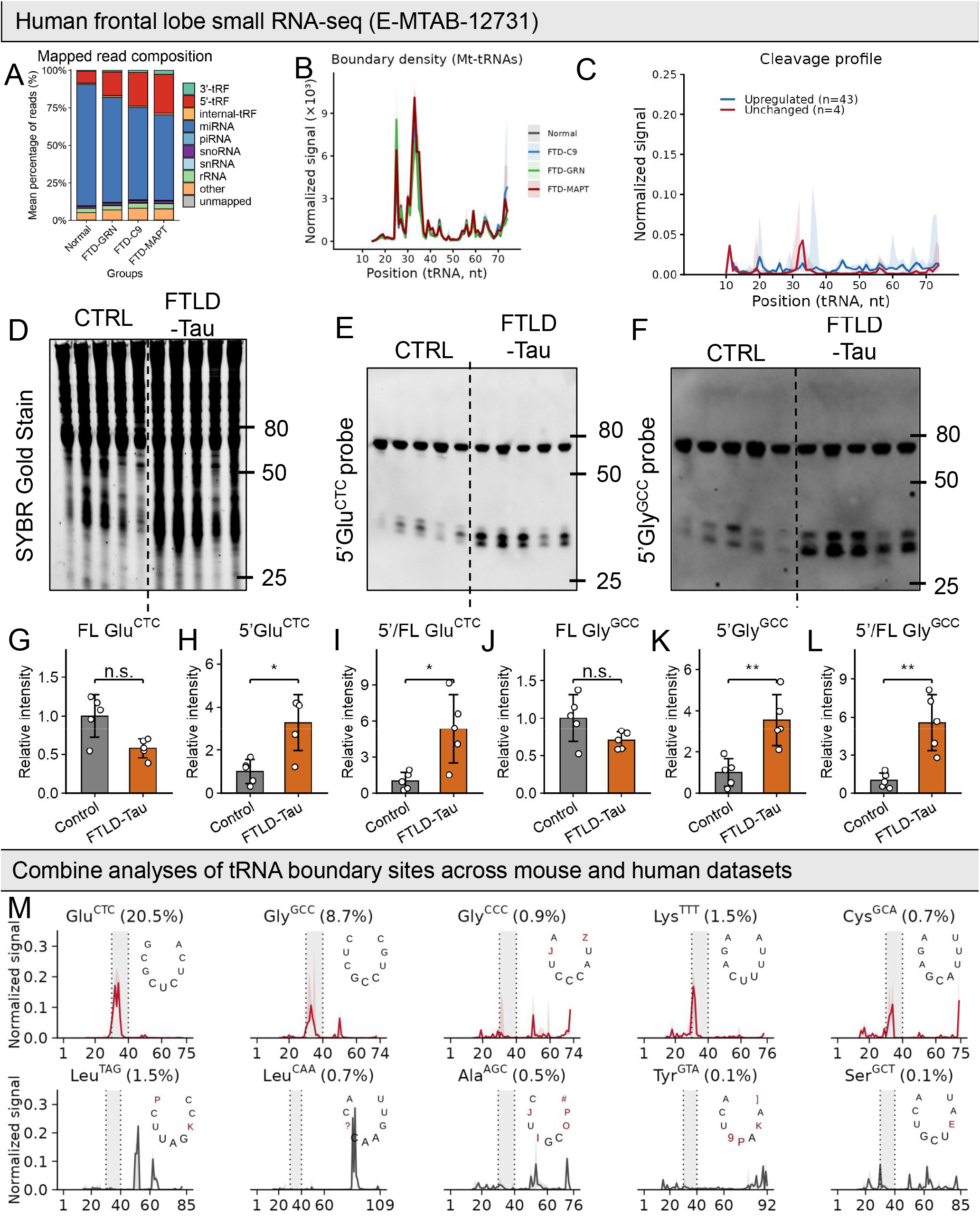
Characterization of tRFs dysregulated in chronic and acute brain disease. **(A-C)** Human frontal lobe small RNA-seq data (E-MTAB-12731; n = 8-13 donors per group). **(A)** Mean percentage of reads mapped to each RNA biotype across FTD subtypes (C9orf72, GRN, MAPT) and normal controls, ordered from lowest to highest total tRF content. **(B)** Boundary density for mitochondrial tRNAs. Major peaks are preserved across conditions, with no differences between control and disease groups. **(C)** Fragment boundary density across the tRNA body (5′ to 3′) for tRNA families unchanged (left) or upregulated (right) in FTD-MAPT relative to normal controls. Lines represent median DESeq2-normalized signal; shaded regions indicate IQR. **(D-F)** SYBR Gold gel and Northern blot of human cortex samples from normal controls and FTD-MAPT donors, probed for 5′Glu^CTC^ and 5′Gly^GCC^. Dashed lines demarcate control and FTD-MAPT groups (n = 2 male, 3 female donors per group). **(G-L)** Quantification of northern blot band intensities for 5′Glu^CTC^ (G-I) and 5′Gly^GCC^ (J-L), shown as relative intensity for the full-length tRNA band (G, J), the 5’ tRF band (H, K), and the 5’ tRF-to-full-length tRNA ratio (I, L). Statistical significance was determined using an unpaired t-test. *p < 0.05; **p < 0.01; n.s., not significant. **(M)** Median DESeq2-normalized signal across tRNA coordinates for selected tRNA families, averaged across seven datasets (two whole brain, three brain regions, FTD, and TBI), with shading indicating IQR. The dashed window (30-40 nt) marks the anticodon loop. Red traces denote tRF families consistently elevated across aging and neurological disease datasets, whereas grey traces indicate families with stable signal. Percentage values represent estimated median abundance within the total nuclear and mitochondrial tRF pool (sum of all three tRF classes). Anticodon loop positions are annotated based on MODOMICS human tRNA references. Custom symbols denote modified nucleotides: J = Um (O2′-methyluridine 5′-monophosphate); Z = Ym (2′-O-methylpseudouridine); ? = tm5s^2^U (5-taurinomethyl-2-thiouridine) and m5C (5-methylcytidine); P = Y (pseudouridine); K = m^1^G (1-methylguanosine 5′-monophosphate); I = I (inosine); O = m^1^I (1-methylinosine); # = Gm (2′-O-methylguanosine); 9 = galQ (galactosyl-queuosine); ] = m^1^Y (1-methylpseudouridine); and E = m6t6A (N6-methyl-N6-threonylcarbamoyladenosine 5′-monophosphate).

## METHODS

### Small RNA-seq data processing

For public dataset analyses, raw sequencing data were obtained from the Sequence Read Archive (Sup. Table 1) and trimmed to remove adapter sequences using parameters specific to each library construction protocol. Following adapter trimming and removal of contaminant sequences (UniVec), reads were sequentially aligned to reference RNA classes in the following order: ribosomal RNA (rRNA), tRNA, miRNA, snoRNA, scaRNA, piRNA, protein-coding mRNA, long noncoding RNA, and miRNA hairpins, with unaligned reads passed to the next class at each step. Multimapped reads were retained and fractionally assigned based on the best alignment. Reference annotations were obtained from RNAcentral (rRNA), GtRNAdb (tRNA), miRBase (miRNAs), piRBase (piRNAs), and Ensembl (snRNA, snoRNA, other ncRNAs, mRNA, and lncRNA). Using this multi-step alignment workflow, an average of 90-95% of reads were successfully mapped across samples. The complete alignment pipeline is available on GitHub (https://github.com/Krichevsky-Lab/smallRNA_seq_pipeline; DOI: 10.5281/zenodo.20140745).

tRNA-derived reads were classified following the tRAX framework (http://trna.ucsc.edu/tRAX/) based on read position relative to mature tRNA boundaries. Reads mapping within 10 nucleotides of the 5′ end were classified as 5′ tRFs, those within 10 nucleotides of the 3′ end as 3′ tRFs, and reads mapping to internal regions not overlapping either end were classified as internal tRFs. Reads spanning both boundaries were classified as full-length tRNAs. Counts for individual tRF isodecoders were then summed into family-level features, and low-abundance RNA species (<1 cpm in fewer than 20% of samples per organ or brain region) were removed prior to downstream analysis. Raw read count tables for each dataset, prior to isodecoder collapsing and filtering, along with corresponding metadata and analysis code, are deposited on Zenodo (DOI: 10.5281/zenodo.20141089).

### Age association enrichment analysis

Read counts were normalized using DESeq2, with size factors estimated from all detected RNA species within each dataset. Age associations for individual tRF species were assessed using Spearman correlation between DESeq2-normalized fragment abundance and age (in months), computed across all samples within each dataset. P-values were adjusted for multiple comparisons using the Benjamini-Hochberg procedure, and a significance threshold of FDR-adjusted p < 0.05 was applied throughout. The percentage of significantly positively or negatively age-correlated species was calculated per biotype (e.g., 5’ tRF, 3′ tRF, internal tRF, Mt tRF, miRNA) and plotted as a diverging bar chart. For human FTD data, differential expressions between each FTD subtype and neurologically normal controls were assessed using DESeq2, with significance defined as padj < 0.05 and |log2FC| > 0.5.

### RNA class abundance trends

For each RNA biotype (5′ tRF, 3′ tRF, internal tRF, miRNA, and snoRNA),DESeq2-normalized counts were summed across all features within each biotype per sample. Per-sample sums were then normalized to the median value of the youngest age group (2 months), yielding fold change relative to baseline. Median and interquartile range were computed per biotype across samples within each age group and plotted as line graphs with IQR ribbons.

### Principal component analysis

PCA was performed on nuclear-encoded tRF count data (5′, 3′, and internal tRF classes combined) normalized using DESeq2-estimated size factors. Features with a mean normalized count below 5 were excluded prior to dimensionality reduction. Normalized counts were log2-transformed (log2(x + 1)) and scaled across features before applying PCA using prcomp. The top two principal components were retained for visualization, with samples colored by age group.

### Fragment-age correlation heatmap

Spearman correlation coefficients between DESeq2-normalized fragment abundance and age were computed for each tRF isodecoder. Size factors were estimated using and applied to the full count matrix. P-values were adjusted for multiple comparisons using the Benjamini-Hochberg procedure, with significance defined as FDR-adjusted p < 0.05. Results were collapsed to the family level by averaging correlation coefficients across isodecoders within each tRNA family and fragment type combination. The top 15 most abundant nuclear-encoded and top 5 most abundant mitochondrial tRF families were selected for visualization, ranked by summed mean normalized abundance across all fragment types. Correlation coefficients are displayed as a heatmap with tiles bordered in black where any isodecoder within the family reached statistical significance.

### Fragment length distributions

Fragment length distributions were computed from interval-level coverage data for each tRF class (5′, internal, and 3′). Read counts were divided by DESeq2 size factors estimated from the full filtered count matrix. Normalized counts were summed per fragment length per sample, and median and interquartile range were computed across samples within each age group.

### Fragment boundary density

The 3′ end position of each aligned tRF interval was mapped to a relative position along the tRNA body by dividing by the annotated tRNA length. Relative positions were then scaled to pseudo-Sprinzl coordinates by multiplying by 76 and rounding to the nearest integer, yielding positions 1-76 corresponding to the canonical tRNA numbering system. Positions within 2.5% of either end were excluded to avoid boundary artifacts. Normalized counts (divided by DESeq2 size factors) were summed per pseudo-Sprinzl position per sample, and median and interquartile range were computed across samples within each age group. An inset zoom panel showing positions 30-35 nt is included.

### Nucleotide-resolution coverage maps

Per-position read depth was computed from interval-level data, where each row represents a tRNA fragment defined by its start and end coordinates and a read count. Each fragment’s count was assigned to every nucleotide position within its span and summed across all overlapping fragments per position per sample. Coverage values were divided by DESeq2 size factors, then summed across isodecoders within each tRNA family. Median and interquartile range were computed across samples within each age group and plotted for Cys^GCA^, Glu^CTC^, and Gly^GCC^.

### Animals

The research complies with all relevant ethical regulations. Animal experiments were carried out in accordance with the U.S. National Institutes of Health Guide for the Care and Use of Laboratory Animals and approved by the Institutional Animal Care and Use Committee at Brigham and Women’s Hospital (protocol 2016N000113). All mice were male C57BL/6 sourced from Charles River. Five-month-old male mice were purchased directly from Charles River, and aged male mice (24 months and older) were generously provided by the National Institute of Aging’s Division of Aging Biology (Charles River colony). All animals were allowed to habituate for at least one week following delivery before euthanasia by CO_2_ inhalation followed by cervical dislocation. Where indicated, brains were dissected into regions prior to freezing. Brains and peripheral tissues were flash-frozen in a methanol/dry ice slurry and stored at -80°C until use.

### Human brain tissue

Frozen human post-mortem brain specimens were obtained from the Massachusetts Alzheimer’s Disease Research Center (ADRC) (grant number: 1P30AG062421-01) and used in accordance with the policies of Brigham and Women’s Hospital’s Institutional Review Board (#2020P002649/BWH). Detailed neuropathological and demographic information is provided in Sup. Table 2.

### RNA Isolation

Total RNA was isolated from snap-frozen brain tissue and peripheral organs using TRIzol (Thermo Fisher Scientific) according to the manufacturer’s instructions. Tissue was homogenized by sequential passage through 16-gauge and 22-gauge needles prior to extraction. RNA was eluted in nuclease-free water, and quantity and quality were assessed by Thermo Scientific Nanodrop 1000 Spectrophotometer.

### Northern blotting and SYBR Gold staining

Total RNA was mixed with 2X RNA loading dye (NEB), denatured at 65°C for 10 minutes, and separated on 10% TBE-urea polyacrylamide gels (Thermo Fisher Scientific) in 0.5X TBE buffer at 200V for 1 hour. Gels were stained with 1x SYBR Gold (Thermo Fisher Scientific) for 10 minutes and imaged using the iBright imaging system prior to transfer. RNA was then transferred onto positively charged nylon membranes (GeneScreen Plus) by wet transfer (1.5 hours, 20V, room temperature) in 0.5x TAE buffer. Membranes were cross-linked by UV irradiation (150 mJ) and hybridized overnight with rotation at 40°C with digoxigenin (DIG)-labeled DNA probes in DIG Easy Hyb solution (Roche). After low stringency washes (twice with 2x SSC/0.1% SDS at room temperature) and a high stringency wash (once with 1x SSC/0.1% SDS at 40°C), membranes were blocked in blocking reagent (Roche) for 30 minutes, probed with alkaline phosphatase-labeled anti-digoxigenin antibody (Roche) for 30 minutes, and washed three times with TBS-T. Signals were developed using CDP-Star ready-to-use (Roche) and detected using the iBright imaging system. DIG-labeled oligonucleotide probes were synthesized by Integrated DNA Technologies (IDT); probe sequences are listed in Supp. Table 3. Densitometry analyses were performed using Image Studio Lite (LI-COR Biosciences).

### RT-qPCR

Total RNA (50 ng) was reverse transcribed using the miRCURY LNA RT kit (QIAGEN, 339340). The resulting cDNA was diluted 80-fold and used for qPCR analysis with the miRCURY LNA SYBR Green PCR kit (QIAGEN, 339345) and miRCURY LNA primers (Supp. Table 4). qPCR was performed on an Applied Biosystems ViiA 7 Real-Time PCR System. Cycling conditions were 95°C for 10 minutes, followed by 50 cycles of 95°C for 15 seconds and 60°C for 1 minute, with a dissociation analysis at the end. Expression levels were normalized to miR-99a-5p and fold changes were calculated using the ΔΔCt method. All critical equipment and commercial kits and reagents are listed in Sup. Table 5.

### Statistical analysis

For validation experiments comparing young and aged mice, statistical significance was assessed using unpaired two-tailed t-tests (n = 6 male mice per group). Results are presented as mean ± SD unless otherwise noted. Significance thresholds are indicated as follows: *p < 0.05, **p < 0.01, ***p < 0.001. All analyses were performed in R.

---

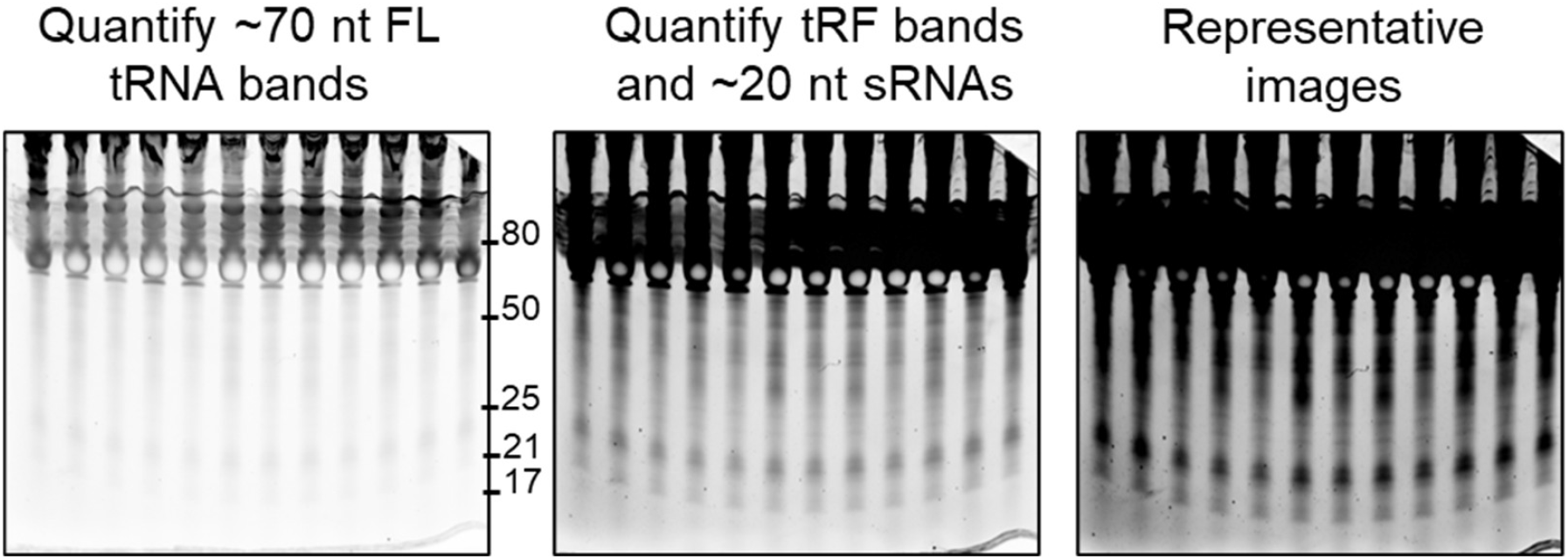

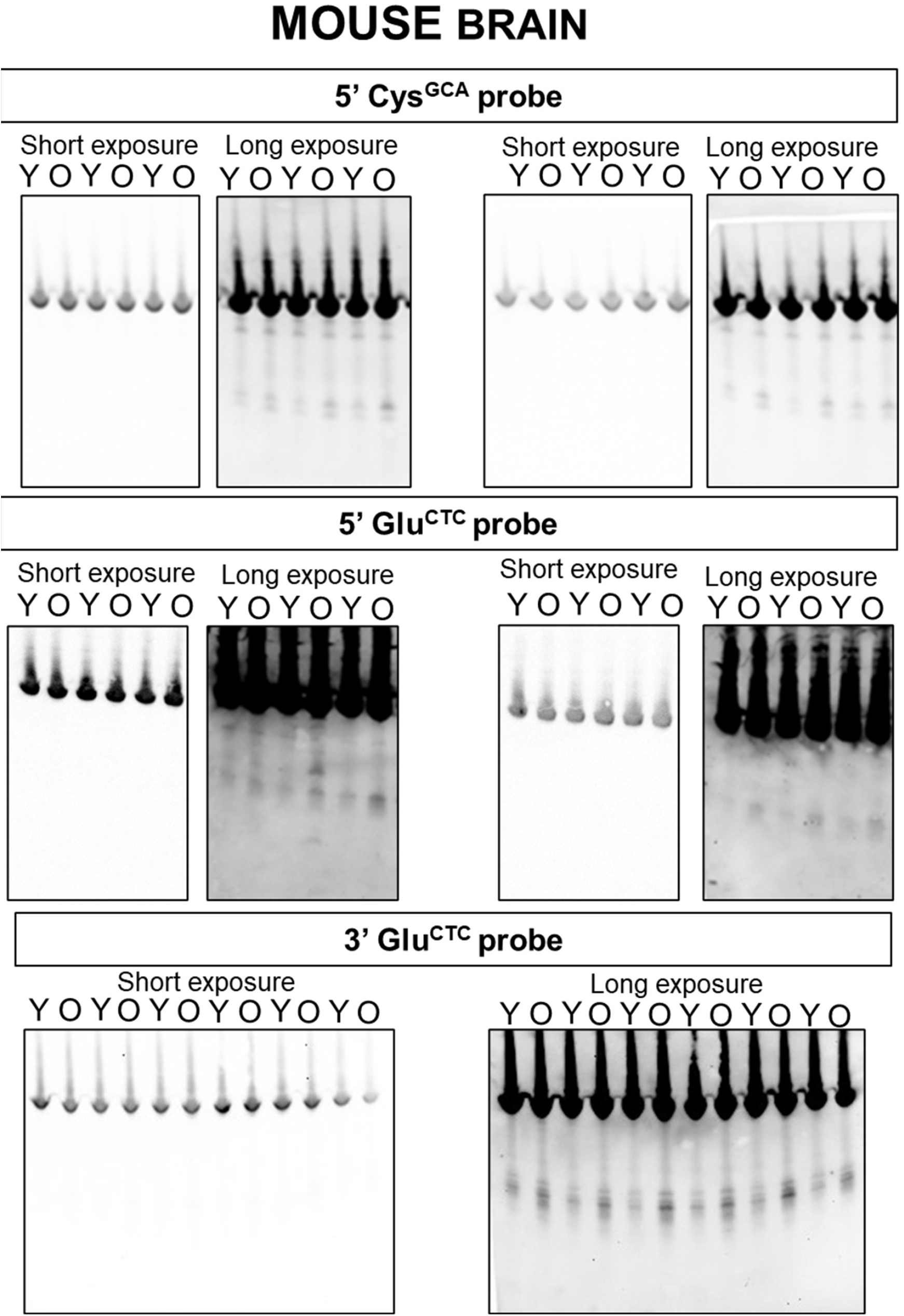

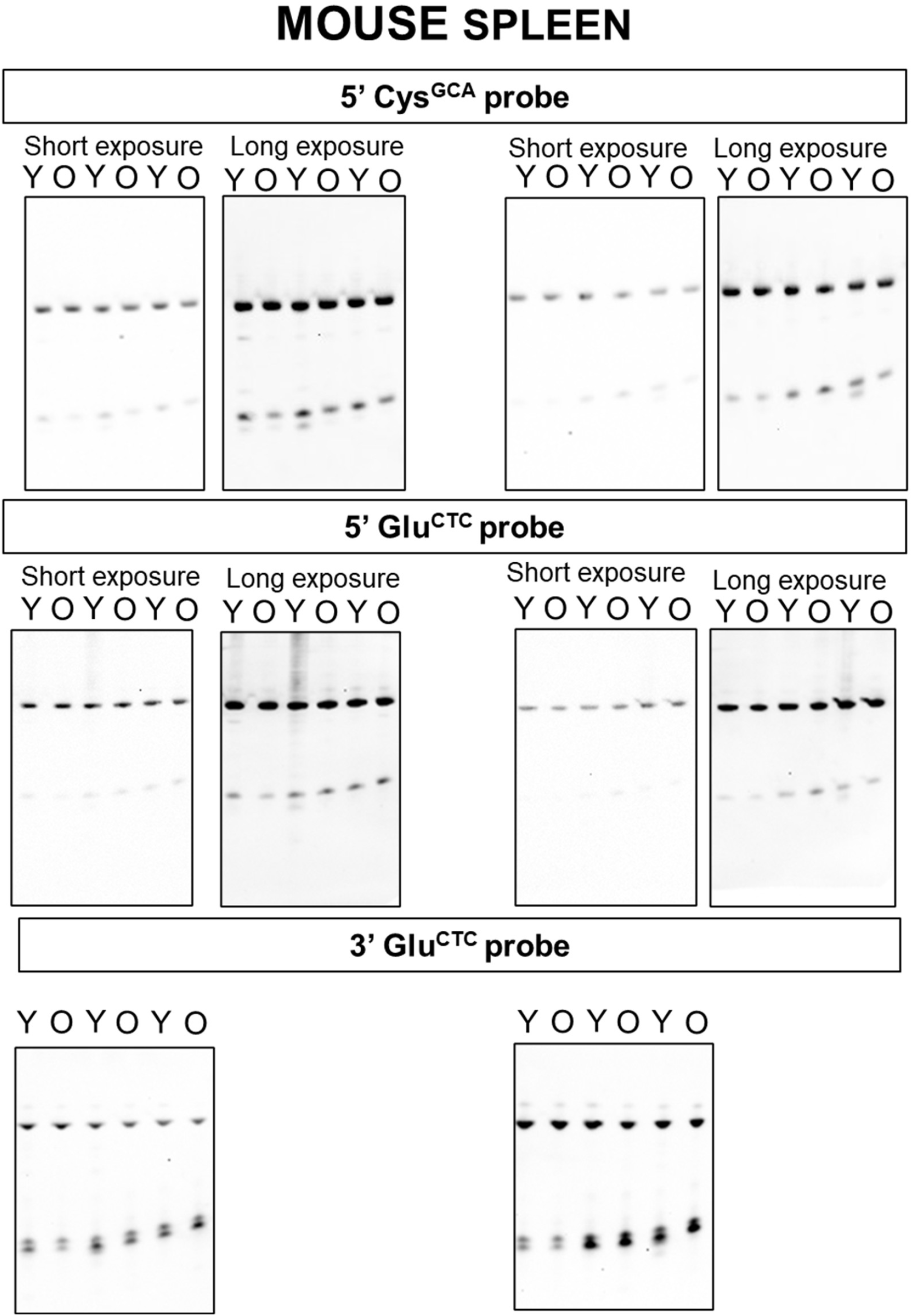

## Notes

https://doi.org/10.5281/zenodo.20141089

https://github.com/Krichevsky-Lab/smallRNA_seq_pipeline

### Competing Interest Statement

The authors have declared no competing interest.

